# Dietary habits of free-ranging banded langur (*Presbytis femoralis*) in a secondary-human modified forest in Johor, Malaysia

**DOI:** 10.1101/2021.03.16.435588

**Authors:** Mohd Faudzir Najmuddin, Hidayah Haris, Noratiqah Norazlimi, Farhani Ruslin, Ikki Matsuda, Badrul Munir Md-Zain, Muhammad Abu Bakar Abdul-Latiff

## Abstract

Banded langurs, *Presbytis femoralis*, are distributed in southern Peninsular Malaysia, i.e., Johor and its borders including Singapore. It has been estimated that there are only < 250 mature individuals of *P. femoralis* in Malaysia and Singapore, and it is currently assessed as Critically Endangered. The dietary information of *P. femoralis* and even other closely related species has rarely been reported. This study, therefore, aimed to describe their dietary habits and discuss interaction between their feeding behaviour and its surrounding. This study was conducted from February to November 2018, with 15 sampling days each month. We collected a total of 186 sighting hours, using a scan sampling method with 10-min intervals, on a five-langur focal group. We have identified 29 species with 47 items consumed by the banded langur, mostly young leaves (51 %) followed by fruits (45 %), and flowers (3.8 %). The study group spent slightly more time consuming non-cultivated plants but relied on cultivated plants for the fruits. Over 75% of fruit feeding involved consuming cultivar plants; in most cases (73%), they ate only the pulp, not the seeds. Since the cultivated plants was planted in human settlement, there is an urgent need to implement conservation measures to untangle the human-langur conflicts for instance, reforestation of a buffer region using non-cultivated plants. There is a potential for building upon our new findings with more detailed investigations, such as more extensive ecological factors influencing the dietary adaptation which would be necessary to support conservation efforts and management decisions of this species.

## BACKGROUND

Non-human primates cover various trophic niches, from being nearly exclusively folivore to being a combination of frugivore, gummivore, insectivore and omnivore. Colobine monkeys, which include ten genera throughout Asia and Africa (Mittermeier et al. 2013), differ from all other primates by having a foregut-fermentation digestive system that enables them to better exploit a folivore diet than other primate taxa (Chivers 1994, Matsuda et al. 2019). However, contrary to earlier assumptions that colobines are almost exclusively folivores, considerable variation in their diet has been reported (Fashing 2011, Kirkpatrick 2011). Indeed, a higher reliance on fruits and/or seeds as food during periods of high fruit availability has been reported in various Asian colobines, e.g. *Presbytis percura* (Megantara 1989); *P. melalophos* (Davies et al. 1988); *P. potenziani* (Hadi et al. 2011); *P. rubicunda* (Davies 1991, Ehlers Smith et al. 2013, Hanya and Bernard 2012); *P. thomasi* (Gurmaya 1986); *Trachypithecus obscurus* (Ruslin et al. 2019); *Semnopithecus vetulus* (Dela 2007, Hladik 1977); *Pygathrix nemaeus* (Phiapalath et al. 2011, Ulibarri 2013); *Rhinopithecus roxellana* (Guo et al. 2007, Yiming 2006, Zhao et al. 2015) and *Nasalis larvatus* (Matsuda et al. 2009, Yeager 1989). Remarkably, colobines living in human-modified habitats like forests with a mosaic of village gardens and rubber plantations (Dela 2011) or urban and agro-forested areas and forest fragments (Ruslin et al. 2019) would tend to consume more fruits than reported in more natural habitats. If the ultimate goal for feeding ecology is to understand the factors affecting these variations across the order primates, we need a detailed description of feeding habits in as many species as possible.

The banded langur, *Presbytis femoralis*, is a member of the subfamily Colobinae found on the Malay Peninsula and Sumatra and was originally classified as three subspecies (Ang et al. 2020b, Nijman 2020, Roos et al. 2014): *Presbytis f. femoralis* (listed as Vulnerable on the IUCN Red List), *Presbytis f. percura* (Data Deficient) and *Presbytis f. robinsoni* (Near Threatened). Abdul-Latiff et al. (2019a) hypothesised using a molecular phylogeny approach analysing Asian colobine DNA sequences that these three subspecies classifications for *P. femoralis* are not suitable because each taxonomic group exhibited separation at the species level in the D-*loop* region of mitochondrial DNA. Due to the unavailability of a DNA sequence from *P. f. percura* in Sumatra, Abdul-Latiff et al. (2019a) were unable to prove this hypothesis and did not elevate these three subspecies to species classifications as *P. percura, P. femoralis* and *P. robinsoni*. They chose instead to differentiate *P. f. femoralis* in Malaysia with the previously listed junior synonym *P. neglectus*. A recent shotgun sequencing of faecal DNA from Asian colobines, however, concludes that those three subspecies of *P. femoralis* are indeed unique species and as such are elevated from their subspecific status (Ang et al. 2020b). As part of this assessment, the conservation status of the three subspecies was also evaluated. *Presbytis f. femoralis* (here after *P. femoralis*), the species distributed in southern Peninsular Malaysia, i.e. Johor and its borders (Pahang state), including Singapore, has been assessed as Critically Endangered (Ang et al. 2020a) as a result. Indeed, it has been estimated that there are less than 250 mature individuals of *P. femoralis* in Malaysia and Singapore (Abdul-Latiff et al. 2019a, Ang et al. 2020b).

The dietary habits of *P. femoralis* and other closely related species have rarely been reported. Notable exceptions are the studies conducted by Ang (2010) and Srivathsan et al. (2016) which describe the plants consumed by *P. femoralis* inhabiting the largest nature reserve in Singapore (53 identified plant species) and by Najmuddin et al. (2019a) in the human-modified forest patch in Johor (27 plant species), respectively. The only long-term study on feeding habits with over 1,500 observation hours found a total of 136 plant species were consumed by the closely related species (or subspecies), *P. percura* (or *Presbytis f. percura*), living in Sumatra, Indonesia (Megantara 1989). Thus, there are very few studies on the feeding ecology of *P. femoralis* even though feeding is one of the most basic aspects of primate ecology that can provide useful information to contribute to the effective conservation management of their habitats.

In studying the ecology of *P. femoralis*, the major challenge in previous work has been a lack of behavioural observations due to both their shy and indolent dispositions and their habitat preference for secondary and freshwater swamp forests that are difficult and dangerous for observers to access (Ang 2010, Srivathsan et al. 2016). We, however, were able to habituate a group of *P. femoralis* to human observation by selecting a forest consisting of mangrove forest, oil palm plantation and lowland forest with fruit orchards grown by villagers in Kota Tinggi Johor, Malaysia. This habitat allows daytime observations of the monkeys on foot. Consequently, authors of other studies have reported the population distribution, activity budget and predation threats of *P. femoralis* at this study site (Abdul-Latiff et al. 2019a, Najmuddin et al. 2020, Najmuddin et al. 2019a, Najmuddin et al. 2019b), though knowledge about the feeding ecology of *P. femoralis* is incomplete. Thus, this study aimed to describe *P. femoralis* dietary habits and discuss the interactions between their feeding behaviour and their surroundings.

## MATERIALS AND METHODS

Observations were conducted from February 2018 to November 2018 in a forest along the Johor River near Johor Lama Village, Kota Tinggi, in Johor, Malaysia (104°00’E, 1°35’N). The study area included mixed vegetation consisting of mangrove forest, oil palm plantation and lowland forest, along with fruit orchards grown by villagers. *P. femoralis* is commonly found in the area with long-tailed macaques (*Macaca fascicularis*) and dusky langurs (*Trachypithecus obscurus*). The mean monthly temperature ranged from approximately 29□ to 3□ and the total precipitation at the site from January 2018 to December 2018 was 2,350 mm (Amin and Othman 2018). The focal group for this study was an all-male *P. femoralis* group well habituated to observers and with easily identifiable individuals. The study group initially consisted of five individuals (from March to June), but was reduced to four (July–November) probably due to the immigration of a male. There was also a one-male-multifemale group consisting of 13 individuals (1 adult male, 5 adult females, 4 sub-adults and 3 infants) in the area, but the all-male group was chosen as the focal group due to the reliability of sightings and higher habituation levels.

We collected behavioural data from the focal group members continuously during our observation period, which lasted from 07:00 (or the time when the group was found) until 19:00 (or the time when the group was lost). We conducted scan sampling at 10-min intervals and recorded the activity (feeding, resting, moving, others) of all visible individuals in the focal group. Feeding was defined as foraging, manipulating and ingesting food material. Resting was defined by a lack of movement from a single place and included activities such as sleeping, grooming and sitting. Moving was defined by going from location to another by any means (i.e. jumping, crawling, walking or climbing). We categorised other activities, such as vocalising or playing, as ‘others’. We also recorded which part of the plant was consumed during feeding sessions and collected plant samples for later species identification. Additionally, we defined whether consumed plant species were cultivated plants as described by the Federal Agricultural Marketing Authority of Malaysia (FAMA 2013).

## RESULTS

We collected data for 41 days during the study period. The total observation time was 186.3 hrs (3433 total behavioural scans), and the mean daily observation time was 4.6 ± 0.8 hrs. Throughout the study period, feeding activities accounted for 26.1% ± 17.4% (894 scans) of the total number of daily observations made. The total number of plant species consumed during observations was 29 and included 47 different plant parts (27 genera, 18 families: **Table 1**). The five most consumed species were *Hevea brasiliensis* (13.9%), *Nephelium lappaceum* (13.6%), *Musa acuminata* (10.8%), *Artocarpus heterophyllus* (8.1%) and *Ficus microcarpa* (6.5%). The study group fed primarily on leaves (51.3% ± 36.6% of daily total scans), followed by fruits (44.9% ± 36.0%) and flowers (3.8% ± 15.5%). They consumed mostly young leaves (91.4%) rather than mature ones. The exploitation of ripe fruit pulp and unripe fruit pulp and seeds from both ripe and unripe fruit accounted for 29.5%, 45.6% and 24.9% of fruit feedings, respectively.

**Table 1.**
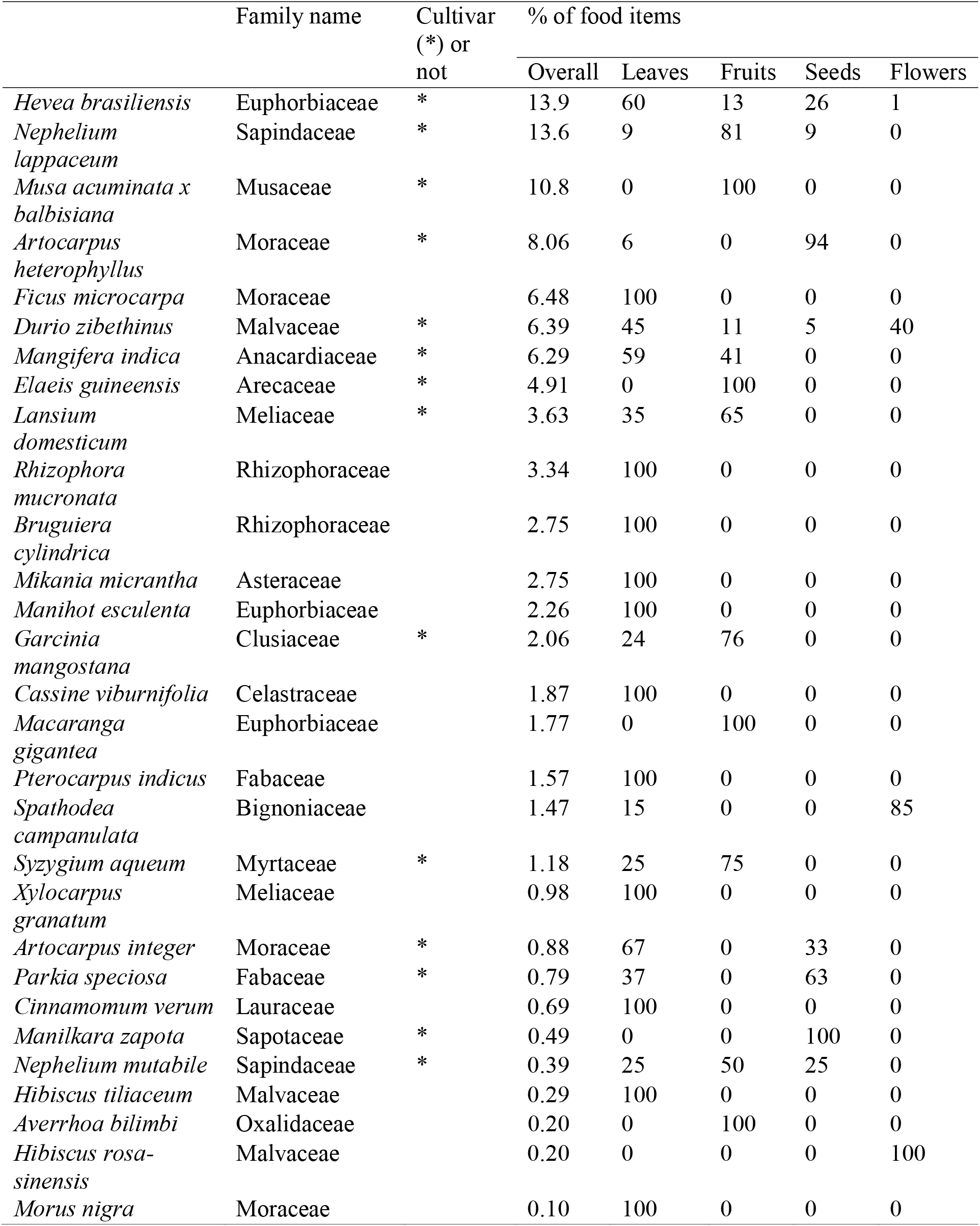
Summary of percentages for food items and parts of each plant species consumed by the study group with categorization if plants are cultivar

Of the 29 consumed plant species, 14 (48.3%) were cultivated plants. The study group spent slightly more time consuming non-cultivated plants (53.8% of total scans), though over 75% of fruit feedings involved cultivated plants. In most cases of fruit consumption (73.2%), only the pulp was eaten and the seeds were untouched (**Fig. 1**). While feeding on the fruit pulp of cultivated plants, the study group often moved from the fruiting trees to safer places further from human presence. This implies that *P. femoralis* may potentially contribute to the seed dispersal of those cultivated plants instead of acting as seed predators. The observed frequency of seed consumption was not related to seed size (**Table 2**); the percentages of seed-eating from each cultivated plant species were not significantly correlated with the maximum seed sizes (Spearman’s rank correlation coefficient: N = 13, rho = 0.38, P = 0.21). This suggests that seed nutrients may be a more important factor for consumption than seed size or amount of pulp.

**Figure 1.**
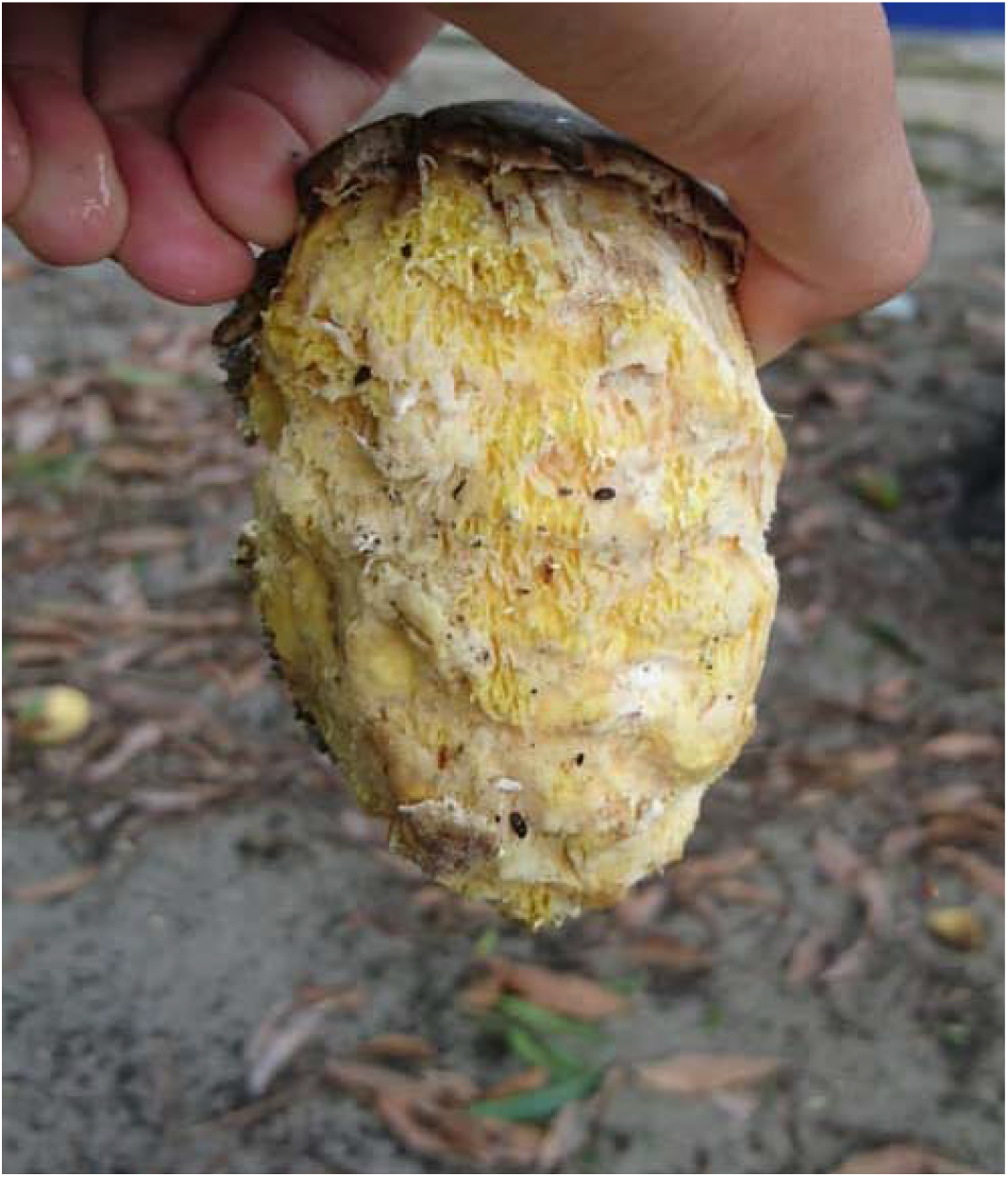
Leftover of a *Mangifera indica* fruit by the focal group of *P. femoralis*; where they mostly consumed only skin and pulp and left the seed out.

**Table 2.** Frequency of seed dispersion and seed predation between 11 cultivated plant species. The size of consumed cultivar plant seeds was also described in case such information is available for published literatures.

| **Cultivated plant species** | **Seed size; Length (L), Wide (W)** | **Seed dispersion frequency** | **Seed predation frequency** | **Reference seed size** |
| --- | --- | --- | --- | --- |
| *N. lappaceum* (Rambutan) | L: 2-3 cm | 112 | 13 | Campos-Arceiz et al. (2012) |
| *E. guineensis* (Kelapa sawit) | W: 1.0-1.8 cm | 43 | 0 | Mohamed Halim et al. (2016) |
| *M. indica* (Mangga) | L: 9-10 cm | 26 | 0 | Campos-Arceiz et al. (2012) |
| *L. domesticum* (Langsat) | L: 1-2 cm | 24 | 0 | Ng (1978) |
| *H. brasiliensis* (Getah) | L: 2-3 cm, W: 1-2 cm | 19 | 37 | Ghani and Wessel (2000) |
| *G. mangostana* (Manggis) | L: 1-2 cm | 16 | 0 | Campos-Arceiz et al. (2012) |
| *S. aqueum* (jambu air) | L: 0.5-1 cm | 9 | 0 | Widodo (2015) |
| *D. zibethinus* (Durian) | L:5-7 cm, W:2-4 cm | 7 | 3 | Berry (1980) |
| *N. mutabile* (Pulasan) | L: 2-3.5 cm, | 2 | 1 | Morton (1987b) |
| *M. zapota* (Ciku) | L: 2 cm | 0 | 2 | Hensleigh and Holaway (1988) |
| *P. speciosa* (Petai) | L: 1.8-2.5 cm | 0 | 5 | Abdullah et al. (2011) |
| *A. heterophyllus* (nangka) | L: 2-4 cm, W:1.25-2 cm | 0 | 74 | Morton (1987a) |
| *A. integer* (cempedak) | L: 3-4 cm | 0 | 4 | Ng (1978) |
| Total percentage |  | 65% | 35% |  |

## DISCUSSION

We describe in this study the feeding behaviour of *P. femoralis* in a disturbed forest in Malaysia. To our knowledge, this is the first report of this species’ feeding ecology, though these results should be considered preliminary due to the small sample size and the limited observation hours. Nonetheless, there is a potential for building upon our findings with more detailed investigations, such as ones exploring more extensive ecological factors influencing dietary adaptations. This information is necessary to support the conservation efforts and management decisions of this species proposed by the immediate change in IUCN status to Critically Endangered by Ang et al. (2020b).

The study group inhabiting the human-modified forest patch, consumed 29 species from 18 plant families, which was lower than that of conspecific inhabiting a species-rich rainforest patch in Singapore (53 species from 33 families), though it was evaluated not only by the direct observation but also metagenomic shotgun sequencing of fecal samples (Srivathsan et al. 2016). Comparing our finding to other studies of Asian colobines, the number of plant species consumed by the study group was particularly low (e.g. *Presbyti spercura*: 136 plant species consumed; *P. melalophos*: 55 species (Davies et al. 1988); *P. potenziani*: 118 species (Hadi et al. 2011); *P. rubicunda*: 64–122 species (Davies 1991, Ehlers Smith et al. 2013, Hanya and Bernard 2012); *Trachypithecus obscurus*: 130 species (Ruslin et al. 2019); *Pygathrix nemaeus*: 79 species (Ulibarri 2013); *Nasalis larvatus*: 188 species (Matsuda et al. 2009). Note, however, that it is impossible to directly compare the number of consumed plant species between this study and others because of differences in sampling methods and habitats. Nonetheless, the low number of plant species consumed in this study may be related to the level of human disturbance occurring in the area. Kg. Johor Lama is filled with human residences, plantations and secondary forests, creating a highly disturbed habitat and, as a result, a loss of forest diversity (e.g. Alroy 2017). This has been shown to decrease gastrointestinal microbial diversity in colobines, which may further be associated with gastrointestinal distress and lead to death (Amato et al. 2016, Clayton et al. 2016, Hayakawa et al. 2018). Due to the extensive habitat loss and forest fragmentation from industrial-scale monoculture plantations in Peninsular Malaysia (Shevade and Loboda 2019), there is an urgent need to evaluate the effects of low dietary diversity on health in *P. femoralis*.

The study group predominantly consumed young leaves (51%); in addition, the percentage of fruits consumed (45%) is comparable to fruit consumption in other Asian colobines, e.g. *T. obscurus*: 40% fruit consumption (Ruslin et al. 2019); *Semnopithecus vetulus*: 60% (Dela 2007); *P. percura*: 58% (Megantara 1989); *P. melalophos*: 50% (Davies et al. 1988); *P. potenziani*: 55% (Hadi et al. 2011); *P. rubicunda*: 84% (Ehlers Smith et al. 2013); *P. thomasi*: 67% (Gurmaya 1986); *Nasalis larvatus*: 40% (Yeager 1989). This frugivore behaviour is also consistent with previous studies of colobines like *T. obscurus* (Ruslin et al. 2019) and *S. vetulus* (Dela 2011) inhabiting human-modified environments. Our study group was observed to consume cultivated plants with fleshy fruits (*Mangifera indica*) and avoid seeds even though it is hypothesised that foregut fermenting primates (i.e. colobines) tend to avoid sugary food sources (i.e. fleshy and ripe fruits). A high-sugar diet can decrease fermentation efficiency or cause acidosis; in extreme cases, it can even result in death among herbivores (Collins and Roberts 1978, Kay and Davies 1994). Our data thus imply that colobine digestive adaptations are more flexible than previously assumed. Supporting this, it has recently been suggested that colobine species with a tripartite stomach such as *T. obscurus* and *S. vetulus* are less susceptible to extreme bouts of malfermentation when fed highly digestible diets like fruits and/or seeds due to a reduced food intake capacity (Hoshino et al. 2021, Matsuda et al. 2019). However, in comparison to *T. obscurus* and *S. vetulus*, the stomach of many foregut fermenting colobine species are poorly understood, including in *P. femoralis*. Thus, investigating forestomach anatomy may be one of the keys to understand their dietary flexibility and adaptability to human-modified habitats.

In Peninsular Malaysia, Afroeurasian monkeys are diverse and human-macaque conflicts, especially *Macaca fascicularis* and *M. nemestrina*, are understood by several places as the result of human settlements neighbouring the natural habitat of macaques, ultimately leading to the labelling of them as ‘pests’ (Abdul-Latiff and Md-Zain 2021, Hambali et al. 2012, Md-Zain et al. 2011). In contrast, reports on human-langur conflicts are generally anecdotal accounts in Peninsular Malaysia and previous data has rarely shown consumption of cultivated plants by langurs. Our study, however, quantitatively demonstrated a reliance on cultivated plants by *P. femoralis*, a species anecdotally believed to be a pest animal in the past. Our study group spent slightly more time consuming non-cultivated plants while at the same time relying on cultivated plants and feeding heavily on their fruits. Since human-langur conflicts may become increasingly serious in Peninsular Malaysia and mirror conflicts seen in other countries (Khatun et al. 2013, Nijman and Nekaris 2010), our study is the first step in understanding suitable habitats for *P. femoralis* and how to protect this species.

Little information is available about crop-raiding by Asian colobines, though a report on crop-raiding by *S. vetulus* in Sri Lanka listed seven high yield crops (coconut, banana, tea, cinnamon, jackfruit, rubber and rice plants) as particularly desirable (Nijman and Nekaris 2010). Of these seven crops, three plants, rubber, banana and jackfruit, were listed as most consumed. However, in some cases of colobines, fruits of cultivated plants dominate their diet even though such fruits are not preferred (Nijman 2012). This suggests that excessive human activity may force *P. femoralis* to consume cultivated fruits despite their natural preferences. Further study assessing not only *P. femoralis* feeding behaviour but also food availability at the study site is necessary to understand the drivers behind their consumption of cultivated plants. This information may lead to a reduction in crop-raiding by colobines through humans adjusting their planting scheme accordingly (Nijman and Nekaris 2010).

Reforestation of our study area with non-commodity plants preferred by *P. femoralis* could serve as a buffer between the forest and other crops and, as a result, reduce their crop-raiding. Promoting ecotourism, and specifically ‘PrimaTourism’ (a term referring to primate-based tourism) (Najmuddin et al. 2019a), at the study site along with broader conservation education programmes may also help increase both income to villagers and awareness for *P. femoralis*. This type of tourism and education has succeeded in protecting other colobines (e.g., Lhota et al. 2019a, Lhota et al. 2019b) and thus provides a useful tool for protecting *P. femoralis* as well. However, a comprehensive framework must be designed for this tourism product, especially on prevention of provisioning by visitors which can trigger human-animal conflict and have negative long-term effects on the species conservation as animals are often over-habituated (e.g., Badiella-Giménez et al. 2021, Mohd-Daut et al. in press, Russon and Wallis 2014), though our preliminary studies show the local people willingness to adopt this species as nature tourism product to generate local economy (Abdul-Latiff et al. 2019b, c).

Lastly, we note that the results of our study should be interpreted with caution because of the limited daily observation time. We successfully observed the langurs when they were both in secondary forest and cultivated area, but when they were in the mangroves or private land (oil palm plantation), we were compelled to stop the observation. Thus, we still cannot deny the effects of such observational bias on dietary diversity and/or their food preference. The langurs may have more diverse diets with less preference to cultivated plants. Until the dietary data of *P. femoralis* based on the full-day observations, have been described, our results must be considered preliminary.

## CONCLUSION

This study provides the first data on the feeding ecology of *P. femoralis* in a human-altered environment in Malaysia. The 29 identified species of plants consumed, consisting of both cultivated and non-cultivated plants, indicates dietary adaptation by *P. femoralis* in a mosaic habitat. Further understanding of feeding ecology, crop-raiding activities and human-langur conflict is critical to formulating a conservation management plan for this species in Malaysia.

## Acknowledgement

This research was supported by Fundamental Research Grant Scheme FRGS/1/2018/WAB13/UTHM/03/2 provided by Ministry of Education Malaysia, and GPPS-UTHM-H552-2019 by Universiti Tun Hussein Onn Malaysia (UTHM). Additionally, it was partly financed by JSPS Core-to-Core Program, Advanced Research Networks (#JPJSCCA20170005 to S. Kohshima) and JSPS KAKENHI (#19H03308 and #19KK0191 to IM). We are grateful to YBhg. Dato’ Abdul Kadir bin Abu Hashim, Director General of the Department of Wildlife and National Parks who provided us with the necessary facilities and assistance. This research was conducted under a research permit (JPHL&TN(IP):100-34/1.24 Jld 8). We are deeply indebted to the Department of Wildlife and National Parks Malaysia for granting permission to carry out this research. Authors acknowledge Ministry of Higher Education Malaysia, Universiti Tun Hussein Onn Malaysia, Universiti Kebangsaan Malaysia and Panz Village for providing necessary funding, facilities and assistance.

We are grateful to Dr. Andie Ang for her comments on the classification and conservation status of the study species.

## Author’s contribution

M.A.B.A. and B.M.M conceptualized the initial idea; M.F.N. performed the behavioural data collection, M.A.B.A arranged the sampling in the field; M.F.N., H.H., N.N., F.R. and I.M. performed and interpreted the analyses; M.F.N., I.M. and M.A.B.A. drafted the manuscript. All authors contributed to the final version of the manuscript.

## Availability of data and materials

Data in support of the findings of this study are available from the corresponding authors by reasonable request.

## Consent for publication

All authors agree to its publication in Zoological Studies.

## Ethics approval consent to participate

Department of Wildlife and National Parks Malaysia that provided with the necessary permission for this research (JPHL&TN(IP):100-34/1.24 Jld 8).

